# Mechanical positive feedback and biochemical negative feedback combine to generate complex contractile oscillations in cytokinesis

**DOI:** 10.1101/2023.12.01.569672

**Authors:** Michael E. Werner, Dylan D. Ray, Coleman Breen, Michael F. Staddon, Florian Jug, Shiladitya Banerjee, Amy Shaub Maddox

## Abstract

Contractile force generation by the cortical actomyosin cytoskeleton is essential for a multitude of biological processes. The actomyosin cortex behaves as an active material that drives local and large-scale shape changes via cytoskeletal remodeling in response to biochemical cues and feedback loops. Cytokinesis is the essential cell division event during which a cortical actomyosin ring generates contractile force to change cell shape and separate two daughter cells. Our recent work with active gel theory predicts that actomyosin systems under the control of a biochemical oscillator and experiencing mechanical strain will exhibit complex spatiotemporal behavior, but cytokinetic contractility was thought to be kinetically simple. To test whether active materials *in vivo* exhibit spatiotemporally complex kinetics, we used 4-dimensional imaging with unprecedented temporal resolution and discovered sections of the cytokinetic cortex undergo periodic phases of acceleration and deceleration. Quantification of ingression speed oscillations revealed wide ranges of oscillation period and amplitude. In the cytokinetic ring, activity of the master regulator RhoA pulsed with a timescale of approximately 20 seconds, shorter than that reported for any other biological context. Contractility oscillated with 20-second periodicity and with much longer periods. A combination of *in vivo* and *in silico* approaches to modify mechanical feedback revealed that the period of contractile oscillation is prolonged as a function of the intensity of mechanical feedback. Effective local ring ingression is characterized by slower speed oscillations, likely due to increased local stresses and therefore mechanical feedback. Fast ingression also occurs where material turnover is high, *in vivo* and *in silico*. We propose that downstream of initiation by pulsed RhoA activity, mechanical positive feedback, including but not limited to material advection, extends the timescale of contractility beyond that of biochemical input and therefore makes it robust to fluctuations in activation. Circumferential propagation of contractility likely allows sustained contractility despite cytoskeletal remodeling necessary to recover from compaction. Our work demonstrates that while biochemical feedback loops afford systems responsiveness and robustness, mechanical feedback must also be considered to describe and understand the behaviors of active materials *in vivo*.

## Introduction

The actomyosin cytoskeleton is an active material that generates contractile forces via the rearrangements and depolymerization of its motor, crosslinker, and polymer components. In non-muscle cells, actomyosin contractility is often activated by the molecular switch RhoA, which, in its active, GTP-bound form, recruits and activates several downstream effectors that recruit, polymerize, crosslink, and remodel the actomyosin cytoskeleton [1]. RhoA is activated in a spatiotemporally specific manner by guanine nucleotide exchange factors (GEFs). Furthermore, RhoA is autocatalytic [2-5]. The RhoA effector Rho-associated kinase (Rho-K) phosphorylates and therefore activates the regulatory light chain of non-muscle myosin II (NMMII). The RhoA effector formin nucleates and promotes elongation of actin filaments (F-actin). Additionally, active RhoA recruits the scaffold protein Anillin that in turn recruits septins. Together, these and other structural elements of the actomyosin cytoskeleton generate forces that maintain or change cell shape [1, 6-10].

In addition to the positive regulation of actomyosin introduced above, contractility is limited by and responsive to changing cues, via time-delayed negative feedback. F-actin promotes accumulation of a RhoA GTPase activating protein (RhoGAP) that inhibits RhoA [2, 11-14]. Via the heterodimeric kinase complex GCK-1/CCM-3, anillin promotes RhoA inhibition by RhoGAP [15]. In addition, RhoA is inhibited downstream of Rho-K [16]. Biochemical circuits that include time-delayed negative feedback loops have been modeled and demonstrated to generate excitability, oscillations, and traveling waves of active RhoA, its cytoskeletal effectors, and contractility [2, 12, 17-21]. Such phenomena have been reported in diverse contexts including cell polarization, mitochondrial homeostasis, and cell division in various fungal and animal cell types [18, 19, 22-26].

Superimposed on positive regulation and biochemical time-delayed negative feedback is mechanical positive feedback, in the form of advective flows that increase local concentrations of all cortical species [27-31]. Early theory work demonstrated that local accumulation of contractile cytoskeleton draws in material centripetally, autocatalytically sustaining and amplifying small, transient local activation [27]. Furthermore, patterning of F-actin assembly via the lateral association of formin with F-actin [32], cooperative binding of F-actin bundling proteins [33-36], biased association of F-actin perpendicular to the curvature of a membrane furrow [37], and mechanoresponsive catch- and catch-slip bonding [38-44] all have the capacity to amplify the activation, accumulation, and rearrangements of the cytoskeleton that comprise contractility.

Though the presence of actomyosin material is often equated with the intensity of contractility, this is an oversimplification. Rather, there exists a complex, non-linear relationship between the abundance of cytoskeletal material and its function (contractility). Too little material is incapable of producing force, while too highly concentrated and crosslinked cytoskeleton cannot contract efficiently; force generation is optimal at intermediate concentrations. This “Goldilocks effect” applies to non-motor crosslinkers and NMMII *in silico, in vitro*, and in several types of animal and fungal cells [45-50]. The Goldilocks effect was demonstrated by comparing contractility of networks with different starting concentrations, but did not take into account how density changes over the course of network contraction. Compaction of a sparse network would boost contractility; compaction of an optimally-contracting network would suppress contractility. The effects of the changes in crosslinker density over time have not been examined.

Cytokinesis is an ancient and essential cell behavior in which localized activation of RhoA leads to the assembly and activation of the actomyosin cytoskeleton (Fig 1A). Cell cycle progression into anaphase and spatial patterning from the anaphase spindle lead to localized activation of RhoA via conserved GEF ECT-2, in what therefore becomes designated as the division plane [51]. As introduced above, active RhoA leads to contractility by recruiting, assembling and activating the actomyosin cytoskeleton [6]. Once thought to be in a different, non-excitable regime during cytokinesis [2, 11], RhoA has recently been demonstrated to undergo time-delayed negative feedback in the division plane [7, 14]. In addition, autocatalytic cortical flows during cytokinesis were modeled to amplify the localized signals of division place specification [27]. At short length scales, material is drawn into contractile foci in some cell types [52-54]. At the whole cell-scale, material flows into the division plane as the cytokinetic ring assembles, contracts, and compacts [28, 29]. Concomitant with concentration via flows, material is lost from the ring over the course of its closure [29, 55-59]. In addition, the “Goldilocks effect” applies to cytokinetic ring starting conditions: cells with lower-or higher-than-optimal levels of crosslinkers furrow more slowly than cells with optimal crosslinker density [45-47, 49, 50, 60]. The co-existence of biochemical negative feedback, mechanical positive feedback, and the non-linear relationship between material density and contractility make their combined effects difficult to predict.

**Figure 1:**
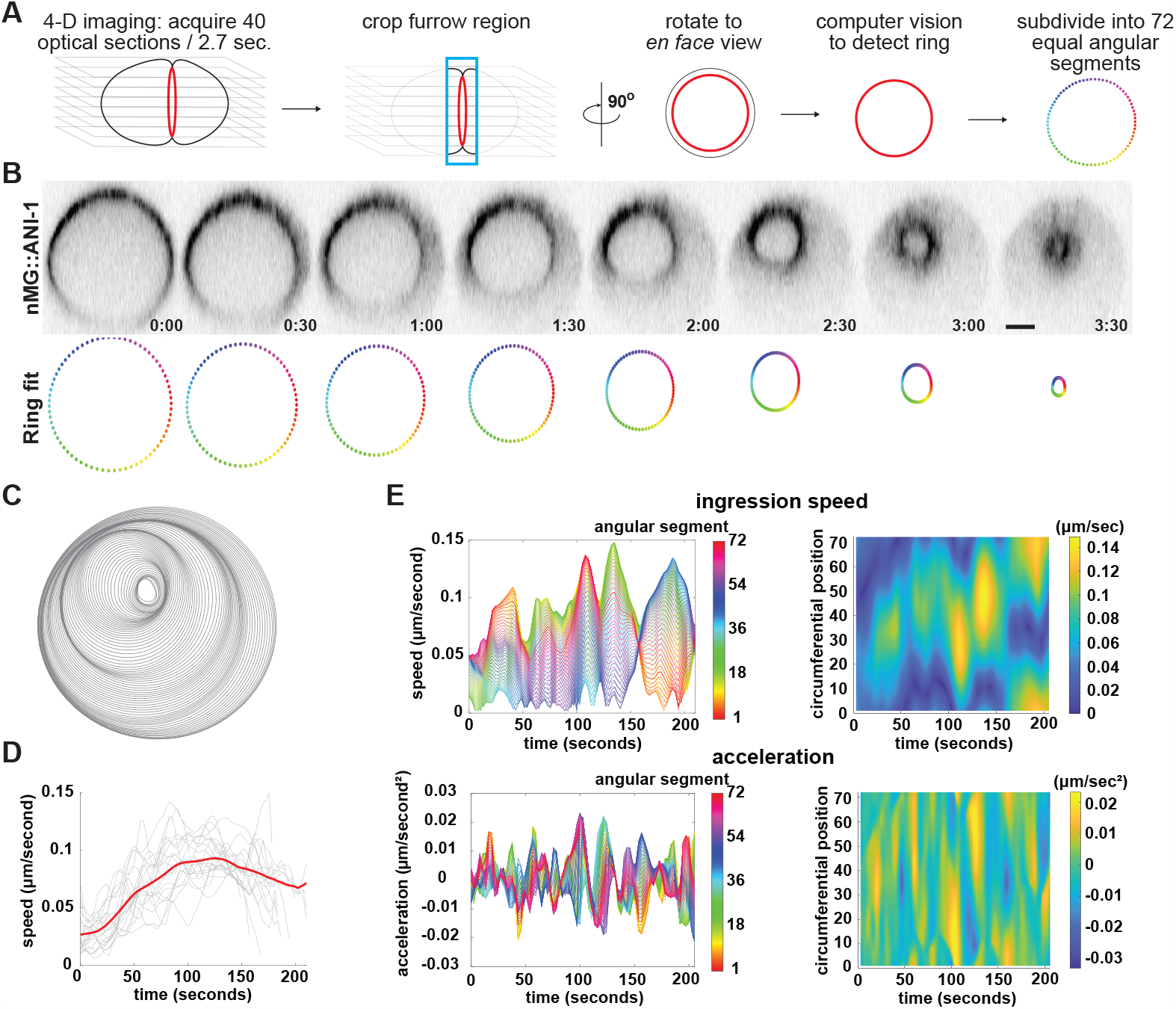
Cytokinetic ring ingression exhibits local oscillation of ingression speed. (A) Schematic representation of the computer vision analysis pipeline to track cytokinetic ring ingression. (B) Representative images of a *C. elegans* zygote cytokinetic ring labelled with mNG:ANI-1 (top) and corresponding segments from fit polygons (bottom). Time labels are minutes:seconds. (C) All fit polygons for a representative embryo. Each polygon represents an individual timepoint. (D) Mean cytokinetic ring ingression speed calculated as the average of all 5 degree angular segments for each timepoint. Grey lines represent speed traces for individual embryos. Red line represents the population mean of all embryos (n=22) smoothed using a rolling average over 5 timepoints. (E) Speed traces of all angular segments of a representative control embryo (top); corresponding acceleration traces (bottom). Left: line plots individual angular segments (color range). Right: Colors represent speed or acceleration; note highlight circumferential and temporal variation of ingression speed.

A recently implemented model of the actomyosin cytoskeleton as an active gel demonstrated that the integration of mechanical positive feedback (*e.g*. due to advective cortical flows) with the well-characterized biochemical feedback loop is sufficient to account for the emergence of complex patterns of contractility with spatial and temporal asymmetries [21]. Because the cytokinetic ring operates under the control of the Rho-actomyosin time-delayed negative feedback loop and experiences cortical flows, we predicted that such complex spatiotemporal patterns occur in cytokinesis. However, the cytokinetic ring has been thought to simply operate at a maximum, and constant, speed [57, 59]. We previously showed that rings in several animal cell types undergo acceleration and deceleration, and exhibit spatial heterogeneities (unilateral closure) even in cases without extrinsic mechanical constraints [37, 61-63]. Higher temporal resolution was required to explore the existence of complex contractile patterns. Since the *C. elegans* zygote is mechanically and biochemically isolated within an eggshell, and has a circular cross-section and centered anaphase spindle, it is an ideal model cell for the study of ring-intrinsic contractility.

Here we establish methods to rapidly acquire image z-series of entire *C. elegans* zygotes, and the computer vision strategies required to annotate and analyze the resulting image datasets. We found that cytokinetic contractility undergoes repeated cycles of acceleration and deceleration. Prominent oscillations with short periods appear to preserve the timescale of RhoA activation pulses, which have a unique timescale in this context. *In vivo* and *in silico*, the period of contractile oscillations was prolonged by the inclusion of mechanical positive feedback. Our work suggests that speed oscillations represent cycles of compaction and remodeling that allow efficient and continual contractility in the cytokinetic ring. Our findings confirm predictions from theoretical work on the effects of combined negative and positive feedback on biological active materials.

## Results

### The *C. elegans* cytokinetic ring undergoes contractile oscillations

The speed of cytokinetic ring contraction is routinely used as a proxy for ring performance. Speed has been variably reported to be constant or to gradually accelerate and decelerate [57, 59, 61]. By contrast, theory predicts that active gels like the actomyosin cytoskeleton, when subject to positive and negative feedback, undergo complex pulsatory dynamics and waves [21]. To explore whether such dynamics exist in the cytokinetic ring, we sought higher temporal resolution imaging of the entire ring. We imaged *C. elegans* zygotes expressing anillin fluorescently tagged at its endogenous locus (mNG::ANI-1; [64]). To eliminate any artifacts resulting from cell compression [65], embryos were not compressed between an agar pad and the coverslip, but instead rested freely on the coverslip in a sealed chamber containing M9 buffer. Imaging was performed on an inverted resonant scanning confocal microscope; we acquired 40 optical sections separated by 1 µm, every 2.7 seconds. We developed a semi-automated image analysis pipeline that tracks ring closure dynamics over time (Fig.1A and B and supplemental movie 1).

Consistent with previous observations that ring closure is asymmetric in the zygote of *C. elegans* and other nematodes [62, 63, 66], different regions of the ring ingressed at different speeds (Fig. 1C). To capture circumferential variation in ring closure speed, we subdivided the ring into 72 equally spaced 5 degree angular segments and calculated closure speed, as the distance between the position of any given segment at time point t and its position at timepoint t+1, for each segment (Fig 1B and E). Individual ring segments did not ingress at constant speed, nor did they undergo a single gradual acceleration followed by deceleration, but rather underwent repeated alternating phases of acceleration and deceleration (Fig 1E and supplemental movie S1). Speed oscillations were non-uniform around the ring circumference; they varied in amplitude, and speed decelerated in some regions while accelerating in others. In addition, the part of the cytokinetic ring that ingressed the fastest was not fixed in place, but instead occupied a different circumferential position at different times during ring closure and traveled circumferentially (Fig 1C and E). We reason that speed oscillations have not been described previously because ring ingression speed has typically been calculated as change in diameter, which is often time-smoothed, and then averaged across multiple cells; these data treatments indeed obscure speed oscillations (Fig 1D).

A segment of the cytokinetic ring could undergo speed oscillations because the cytoskeleton in that region undergoes cycles of activation and inactivation under the local control of regulatory feedback loops. Alternatively, an assemblage of highly contractile cytoskeleton could travel circumferentially, causing local acceleration and deceleration as it entered and exited, a given circumferential position. Some circumferential displacement of cortical material occurs during early anaphase in *C. elegans* zygotes that are compressed against the coverslip but is drastically reduced when compression is avoided [65, 67-69]. We next tested whether the fastest-ingressing region of the ring travels circumferentially because the ring cytoskeleton is physically moving around the ring circumference. To do so, we monitored the circumferential position of fiducial marks of mNG:ANI-1 in the cytokinetic ring. Local mNG:ANI-1 fluorescence inhomogeneities did not travel circumferentially but remained in approximately the same radial position for as long as they could be observed (Figure S1). This evidence that cytoskeletal material does not move circumferentially during ring closure in non-compressed cells suggests that segments’ speed oscillations and circumferential displacement of the region of the ring ingressing fastest occur because different regions of the ring are more contractile at different points in time. This further supports the idea [57, 70] that local assemblages of cytoskeleton, while highly crosslinked within the ring and to the overlying membrane, respond semi-autonomously to biochemical and mechanical signals.

Our novel observation of contractile speed oscillations could be an artifact of fluorescently tagging ANI-1 at its endogenous locus. To test this possibility, we performed the same analysis on cytokinetic rings visualized with endogenously fluorescently tagged non-muscle myosin (NMY-2::GFP), or Lifeact::mKate2 that labels F-actin. In both cases, we observed a similar oscillatory behavior in ingression speed of all ring segments (Supplemental Fig. 2A and data not shown). To determine whether contractile speed oscillations depended on asymmetric ring closure, we performed the same analysis on embryos depleted of either UNC-59 (septin) or ANI-1 (anillin) by RNAi, conditions that strongly reduce ring asymmetry [62]. While cytokinetic rings in cells depleted of septin or anillin indeed closed symmetrically, individual segments still exhibited speed oscillations similar to those observed in asymmetrically closing rings (Figure S2B and data not shown).

Differences in contractile kinetics and molecular requirements exist among blastomeres present in early *C. elegans* embryo [71, 72], so we tested whether cytokinetic speed oscillations are a unique feature of the zygote or occur in embryonic cell divisions more generally. We analyzed ring closure dynamics in the two blastomeres formed via zygote division (P1 and AB), the germline precursor present at the 4-cell stage (P2), and various blastomeres of the 16-cell embryo (Supplemental Fig. S2C). In all cases, speed oscillations were observed and thus can occur across a range of initial cell sizes and cell fates, and thus may be an inherent property of embryonic cytokinesis. In sum, the cytokinetic ring undergoes speed oscillations that qualitatively resemble the complex contractile fluctuations of a simulated active gel controlled by not only biochemical negative feedback, but also mechanical positive feedback.

### The cytokinetic ring contracts with a wide range of oscillation periods

Towards the goal of defining the regulation and mechanical underpinnings of contractile oscillations, and to compare this novel phenomenon in control and perturbed cells, we next sought to quantify the mechanical oscillations we observed. To do so, we measured the period and amplitude of the oscillations of individual segments. Qualitatively, our input data appeared to be both amplitude- and frequency-modulated, so to capture variation in period and amplitude over time, we used wavelet synchrosqueeze transform (WSST) analysis that maximizes temporal resolution while also maximizing frequency space resolution [73, 74] (Fig. 2A). We first calculated acceleration to zero-center our data, and fit the results of the WSST to an adaptive non-harmonic model (ANH) [73, 75]. This approach revealed the amplitude and period of several sinusoidal components (major modes) of the input signal sufficient to reconstruct the input signal (>93% RMSE; Figure 2B). We then tabulated the incidence of all observed oscillation periods, adjusting the counts to account for incidence and amplitude of each mode (see Methods) to plot the distribution of oscillation frequencies present during ring ingression of control embryos (Fig. 2C). This analysis revealed that speed oscillations exhibit a wide range of periods. Since cytokinetic contractility is thought to be under the control of a single biochemical circuit, the coexistence of multiple oscillation periods suggested the existence of additional, coincident feedback loops.

**Figure 2:**
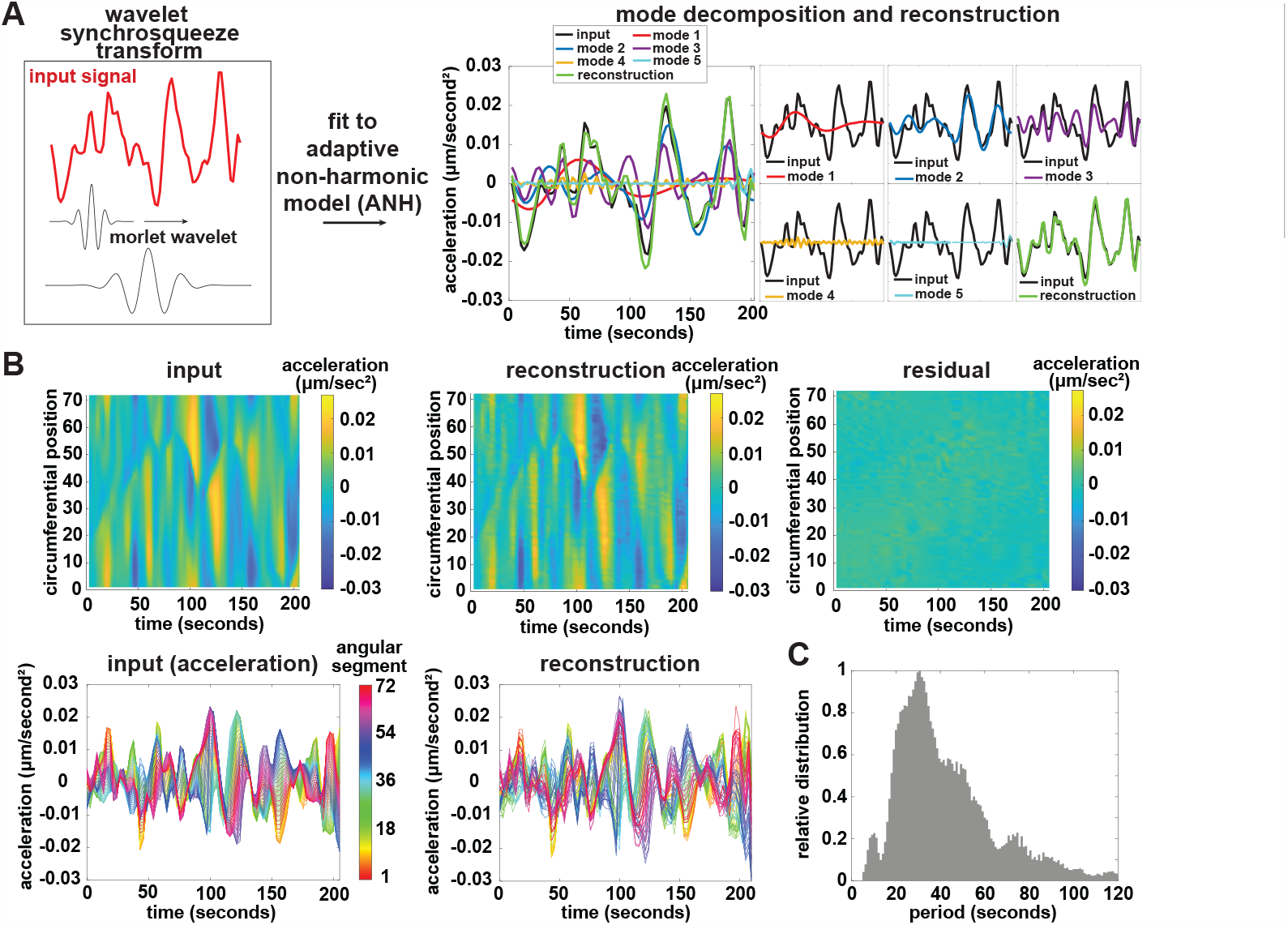
Wavelet analysis quantifies amplitude and period of cytokinetic contractile oscillations. (A) Schematic of wavelet decomposition to determine oscillation modes contributing to speed oscillations. Left: representation of wavelet transform by which the dot product of the input signal (red) and Morlet wavelets of various period (black) was taken. Right: The combination of modes that best recapitulate the input signal are determined by fitting the data to an adaptive non-hormonic model (ANH). Example of a representative analysis of oscillations form an individual input signal (black), that was decomposed into 5 distinct modes (red, blue, purple, yellow, cyan and green) that most accurately reconstruct the input signal (green). (B) Example of WSST and ANH of all 72 segments of a representative embryo showing input signal (left) reconstruction (middle) and residual (right). (C) Histogram showing the adjusted distribution of all oscillation periods from all oscillation modes from control embryos (n=22). Each mode was adjusted by multiplying the amplitude by the inverse of its angular frequency.

### Strain maintains and extends the oscillation period of a simulated active gel and in the cytokinetic ring

Modeled active gel representing the actomyosin cortex exhibited spatially irregular and asynchronous contractile oscillations [21]. The average period of this disordered contractility was 40 seconds when mechanical positive feedback and therefore active contractile stress was minimal, and was longer (60 seconds) when stress was higher [21]. Since contractile oscillations *in silico* qualitatively appeared disordered and heterogeneous, we sought to determine the range of oscillation periods that occur with mechanical feedback. To explain these measurements and to explore how strain and other variables affected the behavior of simulated actomyosin, we used our previously published mathematical model describing the cortex as an active actomyosin gel under the control of a reaction-diffusion model of RhoA activity, both of which were calibrated by *in vivo* measurements from *C. elegans* zygotes (Fig. 3A) [21]. Briefly, this model represents the contractile ring as a one-dimensional domain with periodic boundary conditions, comprising the reaction-diffusion dynamics of an excitable signaling network of RhoA and actomyosin, coupled to the mechanics of the actomyosin cytoskeleton (see Materials and Methods). In this model, RhoA is activated by input stimulus representing the RhoGEF ECT-2 and by autocatalytic positive feedback. An excitable RhoA signaling network generates “actomyosin,” which represents F-actin, active NMMII, and the various cytoskeletal scaffold proteins (including anillin) that are recruited to the cortical cytoskeleton downstream of RhoA. Actomyosin locally inhibits RhoA activity via time delayed negative feedback. Simulating the cortex as a porous Maxwell viscoelastic material allows the calculation of strain generation by actomyosin. Local actomyosin accumulation thus generates contractile stresses that lead to mechanical positive feedback, increasing the local concentrations of both actomyosin and RhoA via advective flow (Fig. 3A).

**Figure 3:**
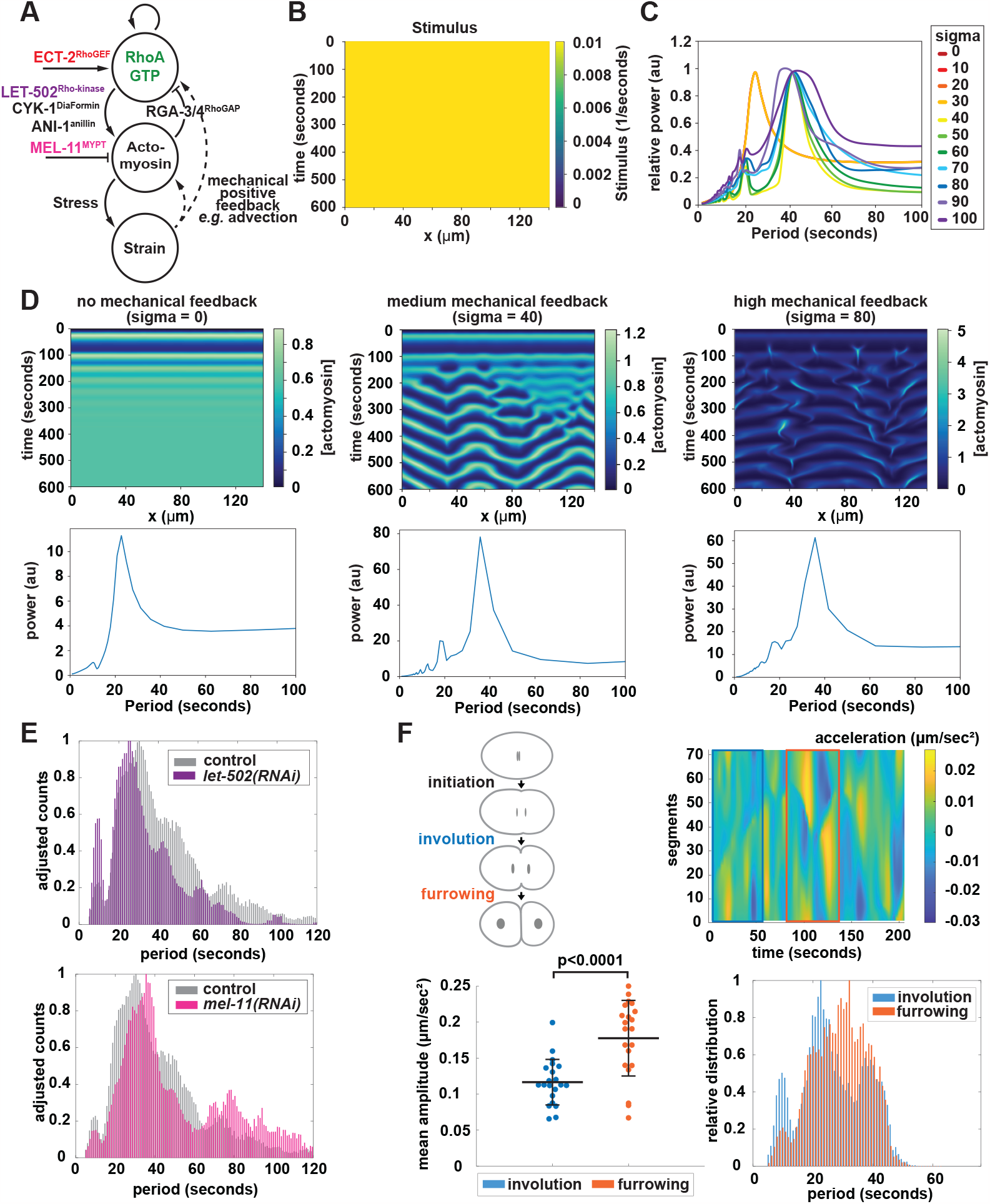
Increased mechanical feedback leads to prolonged oscillation periods *in silico* and *in vivo*. (A) Schematic representation of simulated feedback loops. The point of action of NMMII activator Rho-kinase (LET-502) and the NMMII-binding subunit of myosin phosphatase (MEL-11) are highlighted (B) Kymograph of the simulated continuous stimulus. (C) Plot of frequency spectra of individual Fourier transforms quantifying oscillation periods of simulated actomyosin contractile waves with different levels of strain (sigma). Increasing sigma indicates increases in mechanical feedback. (D) representative kymographs of actomyosin concentrations (top) and corresponding frequency spectra (bottom) in regimes with no mechanical feedback (left), intermediate mechanical feedback (sigma = 40) (middle) and high mechanical feedback (sigma = 80) (right). (E)). Comparison of adjusted period profiles of control and *let-502(RNAi)* embryos (n=10) (middle) and control and *mel-11(RNAi)* embryos (n=10) (right). (F) Schematic of the different stages of cytokinesis in *C. elegans* cytokinesis (top left). Comparison of amplitude (bottom left) and period profiles (bottom right) of speed oscillation during involution (blue) and furrowing (orange). Involution corresponds to the first 55 seconds after furrow initiation while furrowing corresponds to oscillation occurring 81-135 seconds after the onset of furrow ingression (top right).

We first simulated the active gel with a spatially homogeneous stimulus of RhoGEF ECT-2 applied throughout the domain (see Materials and Methods). With sufficiently high stimulus, contractile material accumulated across the one-dimensional space and, after oscillating in abundance several times, reached and remained at a level associated with active contractile stress *σ*_*a*_ [21] (Fig. 3B and D). We then studied the effect of actomyosin-induced contractile stress *σ*_*a*_ on the spatiotemporal patterns of actomyosin. We found that the spatiotemporal distribution of actomyosin material became increasingly complex with increasing *σ*_*a*_, exhibiting irregular pulsatory dynamics (Fig. 3D). We measured the period of the pulsatile contractility using a simple Fourier transform at various levels of active contractile stress (Fig. 3C) and found that as the amount of active stress *σ*_*a*_ was increased, the period of oscillations was higher on average and had a more complex distribution (Fig. 3C and D).

We next sought to test whether tuning active stress *in vivo* led to the same effect of higher average, and more complex, oscillation periods. In the mathematical model, mechanical positive feedback driven by contractile stress takes the form of advective flow, which competes with the diffusion of RhoA and actomyosin. Thus, mechanical feedback is sensitive to changes in actomyosin contractility. To test if contractile oscillations in cytokinetic rings are sensitive to changes in mechanical feedback *in vivo*, we modulated contractile stress experimentally by tuning NMMII activity. To do so we first depleted Rho-kinase (LET-502) by RNAi to reduce NMMII activity [76]; this treatment slowed cytokinetic ring closure (data not shown) [62]. Rho-kinase depleted cells exhibited speed oscillations with lower amplitude and shorter periods (Fig. 3E). We then increased NMMII activity by depleting the NMMII-binding subunit of the NMMII inactivating phosphatase MYPT [76]; this condition shortened the duration of furrowing (data not shown) [76]. In cells depleted of MEL-11, contractile oscillations had longer periods (Fig. 3E). These results supported the conclusion that the long and varied periods of contractile oscillation in control cells reflect the strength of mechanical positive feedback.

Since depletion of conserved, essential proteins may lead to confounding effects, we sought a naturally occurring variation in contractility to test whether that variable was sufficient to explain differences in oscillation periods. In *C. elegans* blastomeres, tension increases over the course of cytokinetic ring closure [77]. We reasoned that this phenomenon also occurs in the zygote, and first compared speed oscillations between early and later furrowing in control embryos. During “involution,” force is exerted across a relatively broad region of the cell and mean ingression speed of the entire cytokinetic ring is relatively low, while during “furrowing” the initially-rounded furrow adopts a planar leading edge and mean ingression speed is higher [78]. Both the amplitude and period of oscillations were significantly higher during furrowing than during the involution phase of cytokinesis (Fig. 3F). These findings aligned with the prediction made by modulating active stress *in silico*, and supported the idea that contractility is highly complex in cytokinesis due to mechanical feedback.

### Contractile oscillations preserve and extend the timescale of RhoA activity pulses

The biochemical oscillator circuit wherein active RhoA elicits actomyosin activation and assembly, which in turn inhibits RhoA, has a characteristic timescale in each context in which it has been examined. In pulses and waves, the abundance of active RhoA and its cytoskeletal effectors rise from and fall back to undetectable levels in 30-120 seconds, depending on the cell type [2, 12, 14, 15, 20, 31]. In the *C. elegans* zygote undergoing polarization during mitotic prophase, the contractile pulse timescale is 30-35 seconds [12, 31]. We reasoned that in cytokinesis, the timescale of RhoA activation-inactivation pulses would fall within the range of contractile oscillation periods we observed.

To determine the timescale of the biochemical circuit driving cytoskeletal activation and assembly, and RhoA inhibition, in cytokinesis, we visualized active RhoA using GFP:LET-502 as a biosensor for RhoA activity (Fig. 4A). We previously established this tagged form of Rho-kinase to be a *bona fide* Rho biosensor that is sensitive to changes in RhoA activity and that localizes to contractile cortical patches in the early zygote [15]. To observe RhoA in a large region of the cortex in a single optical section, we gently compressed embryos between an agarose cushion and the coverslip. Despite the circumferential cortical rotation around the embryo long axis in early anaphase that results from this mounting method [65, 68], transient cortical patches of active RhoA could be observed in early anaphase, primarily in the cell equator. Similar to what is observed during zygote polarization, anaphase patches of RhoA exhibited a time delayed accumulation of NMMII, consistent with the idea that local accumulation of RhoA activity leads to the local recruitment of NMMII (Fig. 4B, supplemental movie 2). We quantified the persistence of cortical patches of RhoA activity and found that the timescale of the RhoA pulse ranged from approximately 15 to 22 seconds, with an average of approximately 18 seconds (Fig. 4C). This timescale is shorter than that of pulses during polarization, suggesting that the biochemical reactions and interactions that drive this conserved biochemical circuit differ between cell cycle phases in *C. elegans* embryos as has been described for the period of cortical actomyosin waves in other animal cell types [12, 14, 20, 31, 79].

**Figure 4:**
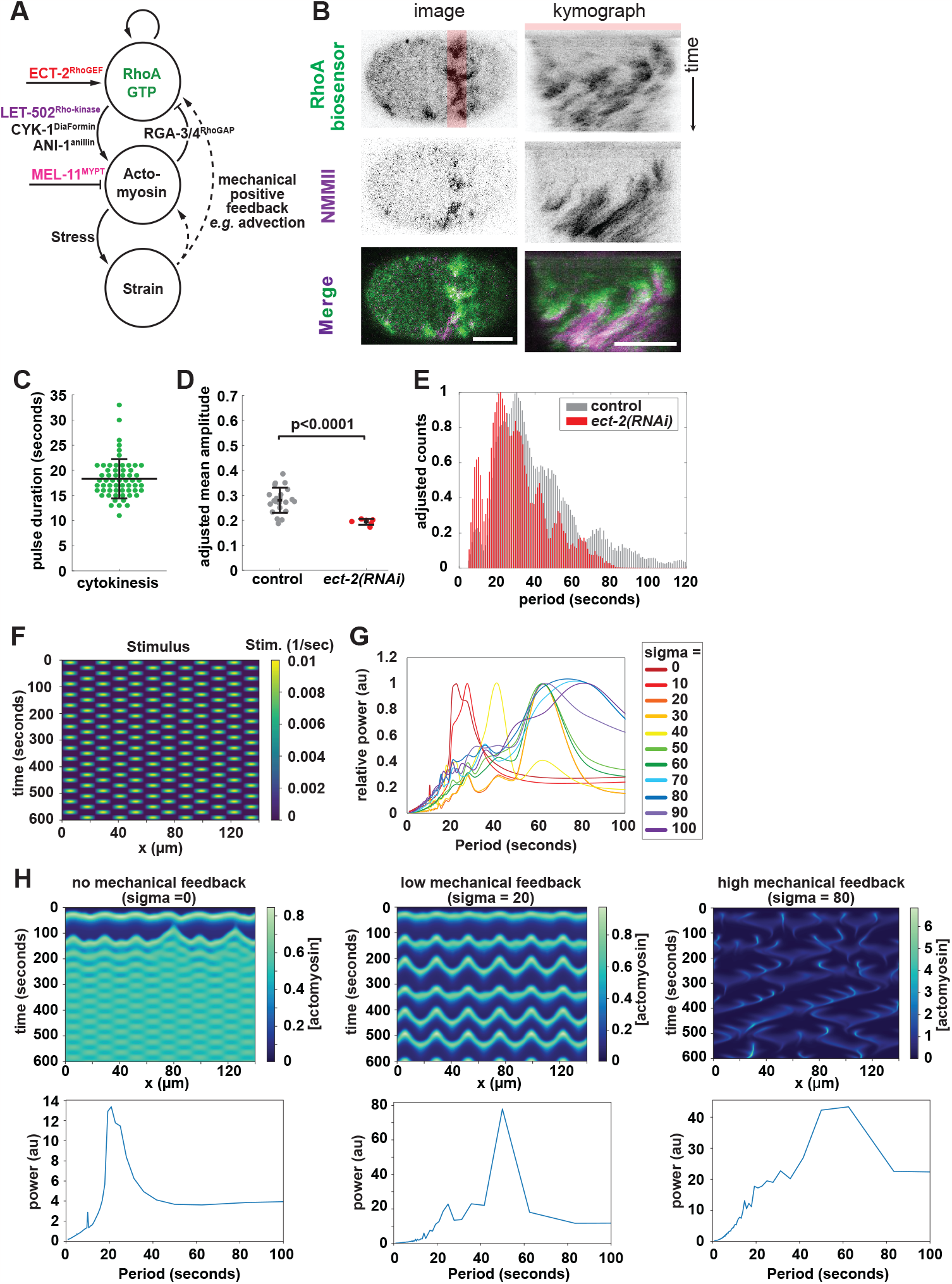
Local speed oscillations are sensitive to changes in the amount of RhoA activity that exhibits pulsed behavior in the cell equator. (A) Schematic representation of simulated feedback loops. Proteins that were targeted for depletion are highlighted with corresponding colored font. (B) *C. elegans* zygote expressing GFP::LET-502 (top, green) used as a RhoA Biosensor and NMY-2::RFP (middle, magenta) and the corresponding merge (bottom). Representative embryo at onset of furrowing (left) and kymograph of the arear highlighted in red (right). (C) Period of GFP-LET-502 pulses in the equator during anaphase (n=51 from 10 embryos). (D) Comparison of adjusted mean amplitude of speed oscillations in control (n = 21) partially ECT-2 depleted embryos with reduced ingression speed (n=6). Individual data points represent mean amplitude of a single embryo. (E) Comparison of adjusted period profiles of control and partial *ect-2(RNAi)* embryos with reduced ingression speed. (F) Kymograph of the simulated pulsatile stimulus. (G) Plot of frequency spectra of individual Fourier transforms quantifying oscillation periods of simulated actomyosin contractile waves with different levels of mechanical feedback (sigma). Increasing sigma indicates increases in mechanical feedback. (H) representative kymographs of actomyosin concentrations (top) and corresponding frequency spectra (bottom) in regimes with no mechanical feedback (left), low mechanical feedback (sigma = 20) (middle) and high mechanical feedback (sigma = 80) (right).

To test how contractile speed oscillations relate to the intensity of biochemical feedback, we decreased RhoA activity in cells by depleting the RhoGEF ECT-2. Lower RhoA activity in ECT-2 depleted cells is evidenced by reduced intensity of the RhoA biosensor [15, 80] and slowly closing cytokinetic rings [3, 81]. We observed that the amplitude of contractile speed oscillations was decreased by ECT-2 depletion (Fig. 4 D-E). The distributions of the periods of contractile speed oscillations were also affected by lowering RhoA activity. Depletion of ECT-2 caused speed oscillations to have shorter periods, on average; shorter periods were over-represented and longer periods occurred less, as compared to the distribution of periods in control cells (Fig. 4E). These findings demonstrating that the periods of contractile oscillations respond to the timescale of RhoA pulses suggest that the former are a function of the latter.

Given the central importance of RhoA activation for driving contractile oscillations, we next considered how *in vivo*, the RhoA-actomyosin circuit becomes activated in distinct foci, not uniformly as we and others initially modeled. To test how localized pulsed activation affects our simulated gel, we adapted the model by applying a locally focused, pulsed stimulus (Fig. 4F, see Materials and Methods). In the absence of mechanical positive feedback (*σ*_*a*_ = 0), the period of the input stimulus was preserved and slightly extended by the accumulation of contractile material. We then turned on the mechanical feedback in the model, and therefore the active contractile stress *σ*_*a*_, and measured the response of the active gel to pulsed RhoA activation. Inclusion of low levels of active contractile stress caused a period doubling of the response, and higher active stress caused even more disordered contractility mimicking the propagating waves and contractile flows reported previously [21]. These highly asynchronous and asymmetric response dynamics had high oscillation periods; longer periods were increasingly represented with increased stress (Fig. 4 G-H) as reported above for active gel with homogenous input stimulus (Fig. 3C and D). These results demonstrate that mechanical feedback can prolong pulsatile contraction beyond the period of a pulsed stimulus. These simulations thus further suggest that *in vivo*, the short-period contractile oscillations we observe arise from the preservation of the timescale of the RhoA circuit and that, in a stress-dependent manner, longer-period oscillations emerge.

### Efficiently contracting cytoskeleton exhibits robust actomyosin accumulation and slow oscillation

The mechanical positive feedback loops implemented in our mathematical model exert their favorable effect by advecting (gathering) both upstream biochemical regulators and downstream cytoskeletal elements [21]. This concept predicts that contractility is optimal where the rate of increase of network components is the highest. We thus next tested whether this occurs *in vivo* by comparing the position of the fastest ingressing segment with the region with the largest instantaneous increase in anillin accumulation. Indeed, the fastest ingressing part of the ring exhibited significantly higher rates of anillin accumulation (Fig. 5 A-B). These results suggest that cytokinetic ring remodeling via advective flows promotes contractility.

**Figure 5:**
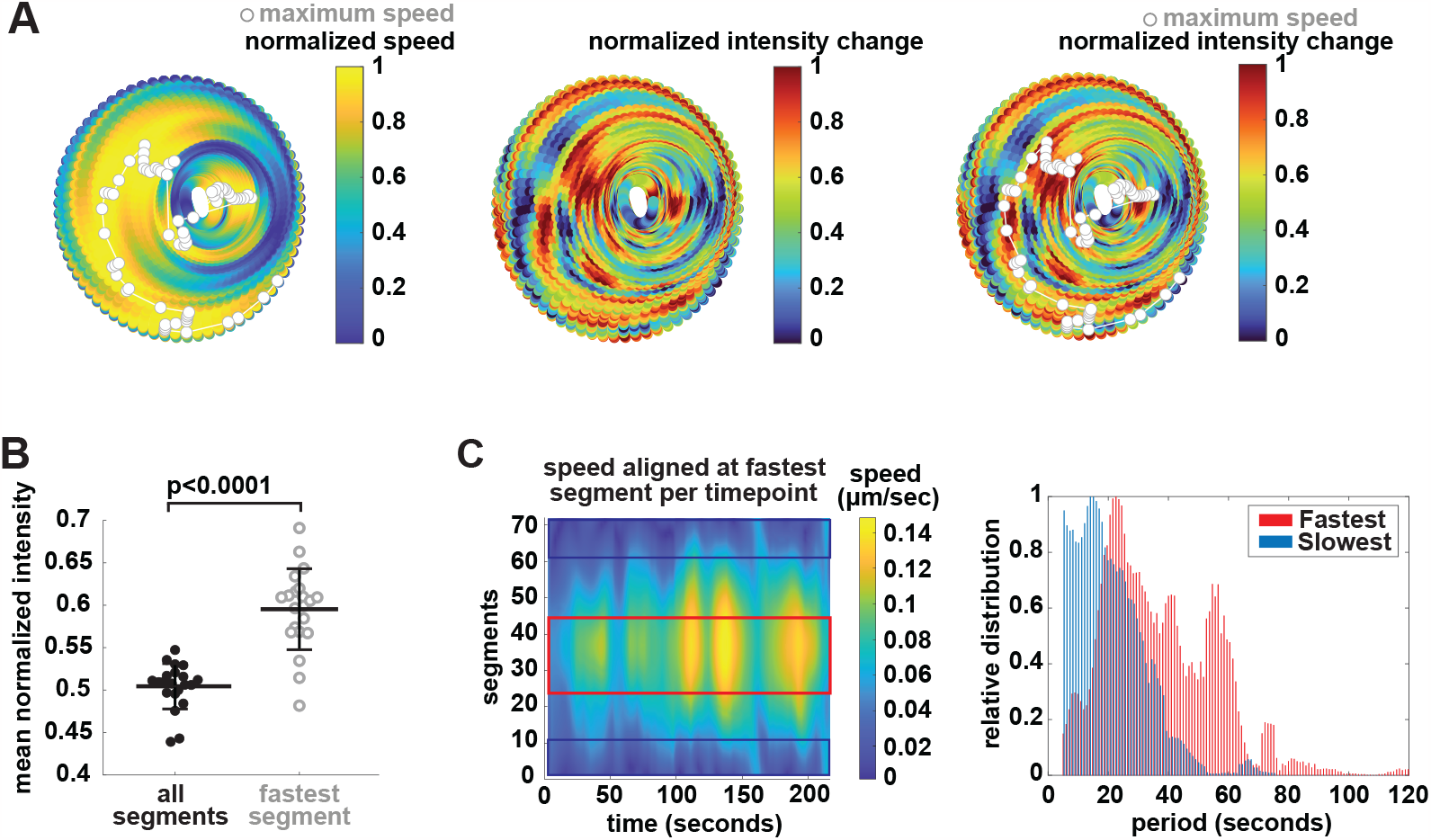
The location of the fastest-ingressing part of the cytokinetic ring correlates with large local increases of ANI-1 density. (A) circular kymographs of measurements from a representative embryo showing local normalized ingression speed (left) local normalized change in mNG-ANI-1 fluorescence intensity (middle) and normalized change in fluorescence intensity overlayed with the position of the fastest traveling segment at each time point (white/grey; right). Each point represents a 5 degree angular segment. Data from each time point were normalized to a range from 0 to 1. (B) Comparison of the mean normalized change in intensity for all segments to the mean normalized change of intensity in the fastest ingressing angular segment only. Each point represents the mean value for all timepoints of an individual embryo (n=21). (C) Period distribution (right) of the 20 fastest (red) and the 20 slowest traveling segments (blue) after aligning segments at the fastest traveling segment at each timepoint (left).

Robust advection and the resulting material compaction thus appear to boost contractility, and predicted to locally generate more stress[45-50]. Therefore, we predicted that local regions of rapid material accumulation and high ingression speed will also experience longer speed oscillations. We next tested whether regions of the ring undergoing fastest ingression exhibited a distinct oscillation signature by comparing the contractile speed oscillations of the 20 fastest moving segments with those of the 20 segments furthest from the fastest-moving segment (which moved slowest). (Fig. 5C). The fast-ingressing segments oscillated with significantly higher period than those around the slowest-moving segment (Fig. 5C). This correlation suggests that speed oscillations with higher amplitude and prolonged period promote contractility. Thus, although contractile speed oscillations arise from the essential and conserved molecular machinery of the RhoA-actomyosin circuit, together with the physical outcomes of cytoskeletal assembly and activity and may thus be inevitable, speed oscillations appear to also endow the cytokinetic ring with a functional advantage via repeating cycles of activation, compaction, and remodeling.

This latter prediction states that the fastest-ingressing part of the ring cannot occupy the same location at all times. This is indeed what occurs in cells, as the fastest-ingressing part of the ring travels circumferentially (Fig. 1C and Fig. S1). To test whether simulated active gel undergoing speed oscillations recapitulates this phenomenon, we modified our one-dimensional model with periodic boundaries so that the domain was allowed to close. This new model consists of a series of active viscoelastic elements, arranged in a ring, that are pulled inward due to actomyosin tension. The magnitude of tension is a function of the local concentration of actomyosin. Rings of active gel elements closed asymmetrically, resembling cytokinetic rings *in vivo* (Fig. 6 A, B). Asymmetric closure was observed despite initially uniform activation of the biochemical and mechanical feedback, as occurs in the *C. elegans* zygote [62]. We also observed the circumferential travel of the region of fastest ingression, as occurs *in vivo* (Fig. 6 C). The circumferential location of highest material accumulation correlated with the region of fastest ingression, as in cells (Fig. 6C, D). In sum, these findings support the conclusion that oscillating between activated and dormant states optimizes the cytoskeletal remodeling that underlies non-muscle contractility.

**Figure 6:**
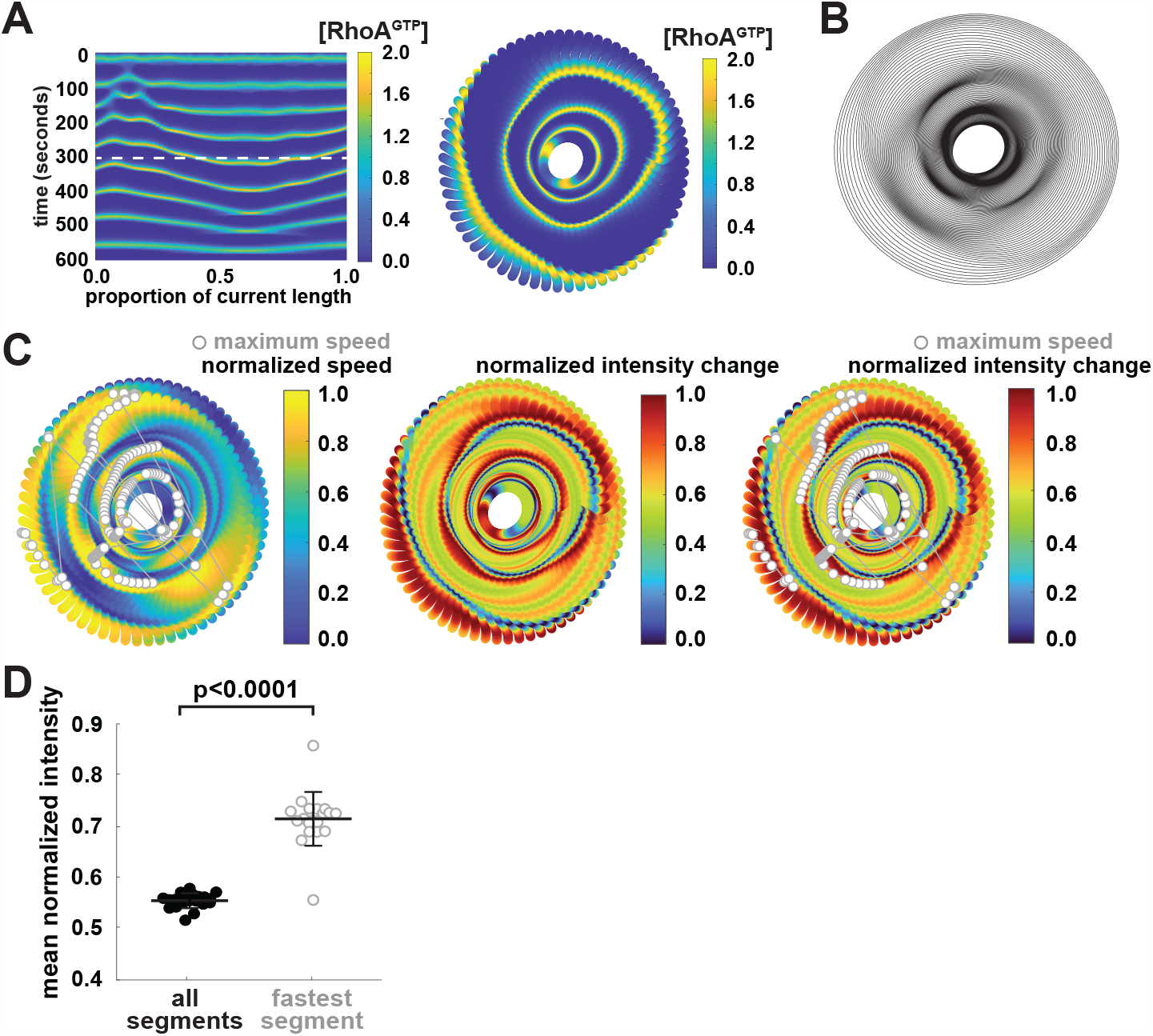
A simulated contractile ring of active gel exhibits physiologically realistic features. Our simulated active gel was allowed to shrink concentrically over time starting 300 seconds after the start of the simulation (white dashed line in (A)). All diagrams except for (A) show simulation results of a representative simulation between 300 and 500 seconds of simulated time. Simulation parameters: strain sigma = 40, stimulus Sr = 0.01, friction in the radial direction gamma_radial = 10, starting ring radius = 15 microns. (A) Kymograph of active RhoA concentration for the entire simulation (left) and Circular Kymograph of active RhoA concentration for timepoints 300-500 (right) (B) Evolution of simulated rings over time, Ring from every other timepoint are shown. (C) Circular kymographs showing local normalized ingression speed with fastest ingressing segment highlighted for each timepoint (light grey) (left), normalized change in local RhoA concentration (values above 0.5 indicate an increase; values below 0.5 indicate a decrease in local RhoA concentration) (middle) and normalized change in local RhoA concentration with the position of fastest ingressing segment highlighted for each time point. (D) Comparison of the mean normalized change in active RhoA concentration for all segments to the mean normalized change of active RhoA concentration in the fastest ingressing angular segment only. Each point represents the mean value for all timepoints between 300 and 500 seconds of an individual simulation (n=20).

## Discussion

Here we report the results of examining a well-studied cell biological subject, the cytokinetic ring of the *C. elegans* zygote, in a new way. By combining unprecedented temporal resolution with observation of the entire ring rather than an optical section, and refraining from excessive temporal smoothing or averaging observations, we discovered a previously unappreciated phenomenon: contractile speed oscillations. Similarly complex spatiotemporal dynamics are exhibited by a simple model of an active gel controlled by a time-delayed negative feedback loop; when mechanical positive feedback is added, the system changes from synchronously pulsatile to exhibiting heterogeneous waves [21]. We showed that *in vivo*, as *in silico*, larger mechanical strain leads to longer-period oscillations. When strain is absent or low, contractile oscillations preserve the period of the input biochemical oscillator, RhoA, which we observed to have a pulse timescale of approximately 20 seconds, shorter than in any other biological context examined. Thus, contractile oscillations emerge from a system of essential, conserved biochemical and structural elements and may be considered inevitable given the time-delayed negative feedback and mechanical positive feedback intrinsic to the contractile cytoskeleton. In addition, we provide evidence that contractile oscillations may endow the cytoskeleton with functional advantages, by counteracting compaction with remodeling.

We further developed our model of the contractile ring as an active gel by allowing the size of the domain to shrink. Previously, we modeled a one-dimensional domain with fixed periodic boundaries [21]. Here, we extend this work by modelling the active gel on an initially circular ring which is able to shrink due to contractile stresses, which can capture the asymmetric contraction rates observed in experiments. This allowed us to pose and test hypotheses about the sufficiency of active gel to recapitulate key characteristics of the cytokinetic ring including ring asymmetry, speed oscillations, and the correlation between contraction speed and material remodeling. Our model was limited to the division plane, omitting the cell poles, which both provide contractile material to the cytokinetic ring [28, 29] and resist division plane forces [58, 82]. Future modeling efforts will be aided by measuring in vivo correlates to the friction within and outside the contractile ring.

Our observations support the hypothesis that cytokinetic rings in animal cells are composed of contractile segments that can contract semi-autonomously [29, 57, 70, 77, 83]. This concept was motivated by observations that furrowing speed in some cell types scales with starting ring size [57, 59], that rings repairing after local laser cutting contract faster than uncut rings [77] and that NMMII organizes in cortical clusters or nodes in *Xenopus* oocytes [70] *C. elegans* embryos [84, 85], human cells [55], and *S. pombe* [86, 87]. Early in *C. elegans* zygote furrowing, when we can reliably assess local cytoskeletal density, the rate of anillin accumulation locally correlates with faster ring ingression speed, suggesting that pulses of RhoA activity drive local contractile oscillations. Such local RhoA pulses lead to the time delayed accumulation of cytoskeletal elements. This leads to the local increase in cortical contractility that initiates a positive mechanical feedback loop recruiting more contractile material from neighboring areas of the cortex via advection [52-54]. Our ability to resolve foci of anillin, a reasonable candidate for a segment marker [12, 52-54, 87], suggests that the *C. elegans* zygote ring comprises ∼24 initially-4-micron segments. By contrast, the effective lengthscale of local contractility may be closer to 18 microns, the hydrodynamic length measured in the *C. elegans* zygote during polarization [88]. Despite the possible autonomy of ring segments, the fact that ingression speed is maximal at a single circumferential site at any timepoint likely relates to the high degree of mechanical coupling among segments via F-actin, cytoskeletal crosslinkers, and scaffolding to the plasma membrane.

While the pulsed nature of the localized activation of RhoA in the equatorial cortex is certainly an important driver of the observed speed oscillation, it is likely not sufficient to fully explain the observed speed oscillations. Indeed, the period of contractile speed oscillations is significantly longer than the period of RhoA activity pulses consistent with the existence of positive mechanical feedback discussed above. Advection driven accumulation of NMMII that can act as both a motor and a crosslinker [47, 89-92] and the non-motor crosslinker anillin will eventually drive the local crosslinker concentration into a regime where mechanical feedback begins inhibits local contractions [48]. For the cytokinetic ring to maintain high overall ingression speed, other contractile segments must become more contractile as the fastest ingressing segment decelerates as evidenced by circumferential traveling of the fastest ingressing segment (Fig. 5A). This local travel may be promoted by a pulse of RhoA activity in a nearby region of the ring initiating a new positive feedback loop. Whether this process is stochastic or influenced by the position of the decelerating ring segment is not clear but since the asymmetry of ring ingression is maintained for longer than individual contraction periods, and since local contractions oscillate, the contractile state of a newly stimulated segment may contribute to whether it will accelerate. The occurrence of speed oscillations in cytokinetic rings of blastomeres at various developmental stages (Supplemental Figure 2C) with varied size and degree of intercellular contacts demonstrates that contractile oscillations are not a phenomenon specific to large, isolated cells such as the *C. elegans* zygote. It will be interesting to quantitatively compare oscillation dynamics among cells of different sizes and developmental fates.

We have stated that the mechanical positive feedback in the model implemented here represents how advection boosts the local concentration of contractile cytoskeleton [21]. While there is ample precedent for the assertion that advective flows act in this fashion [27-29, 31, 52, 88], it is likely that additional mechanical factors contribute to positive feedback. Cortical flows also promote cytoskeletal alignment compatible with contractility [28, 93]. The membrane curvature of the furrow may promote circumferential cytoskeletal alignment [37]. F-actin bundling promotes the association of additional binding proteins with the same filament spacing preference [35, 36]. Extensile strain on F-actin can change the binding of partners [94]. Compaction can bring a network with suboptimal crosslinking density into a regime more effective at contractility [48]. Stress exerted through catch-bonding myosins prolongs their interaction with F-actin [38, 39]. All these phenomena are likely to occur during the remodeling of the non-muscle actomyosin cytoskeleton, and thus may contribute to the contractile oscillations we report. While all these factors occur in response to contractile stress, their differences should be distinguishable via quantitative cell biology. Here, we compared contractile speed with the change in density of one network component, the scaffold protein ANI-1. Future work will be aimed at designing assays for local F-actin bundling, order, and strain to test how these other network remodeling events correspond to contractile speed and its oscillations.

Each of the mechanical positive feedback processes enumerated above has specific molecular mechanisms. The molecules that contribute to these mechanisms have been demonstrated by biophysical assays to bundle, crosslink, mechanosense, and exhibit catch-or catch-slip bonding [1]. Conventional cell biological studies after removing protein function have define proteins as being generally necessary for contractility. Our future work will leverage the measurement of contractile speed oscillations to assign specific physical contributions *in vivo*. For example, proteins that increase the abundance of unbranched F-actin networks at the expense of branched networks could limit mechanical positive feedback by coarsening network heterogeneities [88]. Alternatively, this could drive mechanical positive feedback by promoting filament bundling and the accumulation of F-actin tensile strain. Testing how such perturbations affect contractile speed oscillations will thus offer novel insight into the mechanical contributions of conserved structural and regulatory ring components.

The high spatial and temporal resolution afforded by simulations (using Cytosim or other approaches) may allow us to test how advective flows and other forms of mechanical positive feedback correlate with local contractile dynamics of simulated rings [58, 95, 96]. We envision that contractile oscillations comprise network compaction (acceleration) and remodeling (deceleration). Correlating contraction speed with cytoskeletal advection, compaction, alignment and turnover in silico and in vivo will reveal mechanisms of positive and negative feedback. Regardless of the specific aspects of cytoskeletal remodeling that underlie contractile speed oscillations, this novel phenomenon nonetheless highlights the impacts of coexisting negative and positive feedback loops and provides unique insights into the dynamics of biological active materials such as the non-muscle actomyosin cytoskeleton.

## Acknowledgements

The authors are grateful to Ingrid Daubeschies, Hau-Tieng Wu, Ty Hedrick, and J. Steve Marron for their guidance with signal analysis, Paul S. Maddox and the members of both Maddox labs for valuable discussion, and many other members of our mathematics, physics, and cell biological communities for their insights on this project. This work was supported by the National Institute of Health under award number 1R35GM144238 to ASM and National Science Foundation under award number NSF MCB-2203601 to SB.

## Author contributions

Conceptualization, M.E.W., A.S.M., M.F.S and S.B.; Methodology, M.E.W., A.S.M., D.D.R.,C.B, M.F.S and S.B..; Software, M.E.W, D.D.R, C.B, M.F.S and F.J.; Formal Analysis, M.E.W.; Investigation, M.E.W.; Data Curation, M.E.W, D.D.R and C.B; Writing – Original Draft, M.E.W and A.S.M; Writing – Review & Editing, M.E.W, A.S.M, M.F.S, S.B.; Visualization M.E.W and A.S.M; Funding Acquisition, A.S.M, S.B,

## Material and Methods

### *C. elegans* handling and RNAi

Worm strains (Table 1) were maintained at 20°C as described previously [97]. LP162 was a gift from Dan Dickinson and Bob Goldstein. RNAi was performed by feeding HT115 *E. coli* bacteria expressing double-strand RNA at 20°C as described previously for 24 hours unless specified otherwise [98, 99]. For partial *ect-2(RNAi)*, worms were incubated for 20-22 hours at 20°C and only embryos with at least 20% reduction in overall closure time but still completing cytokinesis were used for analysis.

**Table 1.**
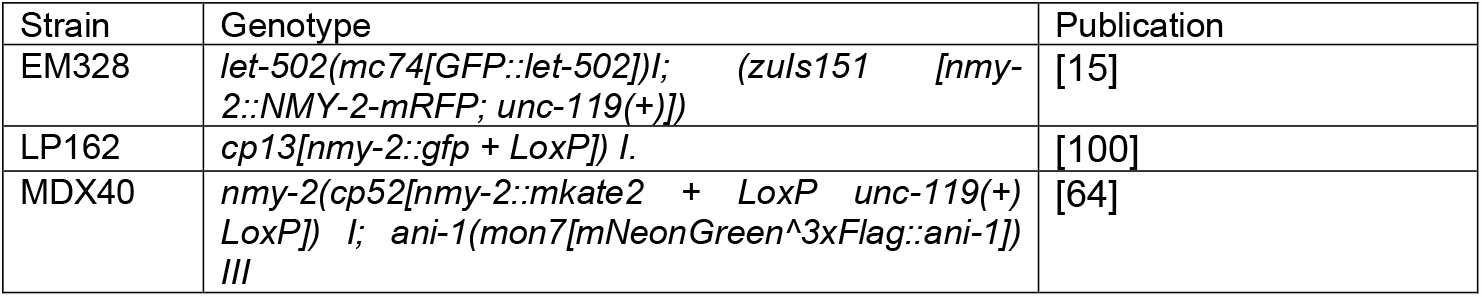

### Live imaging of *C. elegans* embryos

For all experiments except Figure 4 B and C, embryos were dissected from gravid hermaphrodites into a VLAP chamber on a slide containing M9. The chamber was covered with a coverslip and sealed using VLAP. For microscopy, chambers were inverted and embryos were imaged freely resting on the coverslip to avoid compression. For Figure 4B and C gravid hermaphrodites were dissected into M9 on a coverslip that was inverted on a 2% agar pad and sealed with VLAP.

All acquisition was performed on a Nikon A1R microscope body with a Gallium arsenide phosphide photo-multiplier tube (GaAsP PMT) detector using NIS-elements. A 60 × 1.41 NA Nikon Oil Immersion Objective (Figure 4 B and C) or a 60 × 1.27 NA Nikon Water Immersion Objective (all other figures) were used for image acquisition. For figure B and C data were acquired from single Z-sections at the embryonic cortex with a sampling frequency of 1 s using the Galvano point scanner. Acquisition for all other figures were performed using the resonance scanner acquiring 40 confocal sections with 1 μm spacing through the entire embryo every 2.7 s with a PI (Physik Instruments) 100um stage insert Z piezo triggered directly from the A1 confocal controller.

### Analysis pipeline for 4D image acquisition

A 10 µm area surrounding the cytokinetic furrow was selected and an end-on reconstruction was created using the Reslice function in ImageJ. The resulting image stack was average-intensity projected into two dimensions [101-103]. The resulting tiff stack was segmented using a custom Python script that fits a polygon to the cytokinetic ring based on mNeonGreen:ANI-1 fluorescence intensity. The resulting polygon was visually verified and divided into 72 equal angular segments and the coordinates for the center of the segment were recorded for each time point. The Fourier coefficients of the resulting coordinates were spatially smoothed by fitting a 2D harmonic series with 3 harmonics and temporally smoothed using a moving average over 5 time points. A smoothed inward velocity of individual segments was derived using a smooth differentiator function described previously [104]. The resulting velocity and acceleration data was further analyzed using custom Matlab scripts. Mode decomposition of the acceleration input signal was performed using the Matlab wavelet synchrosqueeze transform function (WSST) function using Morlet wavelets as a Kernel and 32 voices per octave [73, 74]. The 25 highest energy “ridges” in the wavelet synchrosqueeze transform matrix that represents the signal were identified using the Wsstridge function in Matlab with a penalty of 2 and 2 frequency bins. Each of these ridges represented a part (or mode) of the signal, which was ultimately reconstructed via addition of the modes. The various mode combinations from the adaptive nonharmonic model were fit to experimental input data. For each cell and spatial segment, we found the representation of that signal as a maximum of 5 simple waves that would reconstruct the input signal with a minimal root-mean-squared error (RMSE). The amplitudes of each mode for the model fit with the lowest RSME was obtained from the Hilbert transform of the corresponding intrinsic mode functions and the mode periods were calculated as the inverse of the frequency output of the WSST. Weighted histograms were created in Matlab by aggregating all mode frequencies from the ANH fit with the lowest RMSE and adjusting the amplitudes by their corresponding angular frequencies and normalizing the data to the frequency bin with the largest amplitude [105].

### Quantification of the period of RhoA pulses

LET-502:GFP foci were identified and tracked using Fiji plug-in TrackMate [15, 106]. Analysis of LET-502 foci was performed on single cortical plane images of the first 2 minutes following anaphase onset in a 10um region surrounding the presumptive site of furrow ingression. Prior to analysis using TrackMate, single cortical plane images were normalized in Fiji. In TrackMate the LoG detector function with an estimated foci diameter of 3 μm and median filter thresholding was used to detect foci and the LAP tracer function with “no gap closing distance” and maximum linking distance of 2 μm was used to track foci. Processing and analysis of foci properties determined using TrackMate were performed using custom scripts in MATLAB. Pulse periods were also scored manually by visually identifying the first and last frame for at least 5 foci for each scored embryo resulting in similar average pulse periods as the automated tracking approach using Trackmate.

### Calculation of change in fluorescent intensity

Using a custom Matlab script, pixels corresponding to the center of the 72 angular segments derived from the segmented polygon described above are identified on the average-intensity projected image stack used for the segmentation. Intensity values for each pixel as well as the 8 pixels surrounding the pixel are extracted and averaged for each segment at each timepoint to obtain a mean intensity value for each segment. Change of intensity for each segment at each timepoint (t) is then determined by subtracting the intensity at timepoint t+1. Intensity values are smoothed by a moving average over 2 timepoints and normalized to the range of intensity changes for each timepoint.

### Quantification of the angle of circumferential travel

The angles between the time axis and the manually assigned vector of displacement in kymographs from intensity or velocity kymographs of the 42 angular segments closest to the coverslip are determined using a custom Matlab script. Resulting absolute angular value distributions are represented in rose plots generated in Matlab. For kymograph analysis intensity and velocity values for individual segments were range normalized for each time points.

### Active gel model of the actomyosin cortex

Active gel model in 1D domain with periodic boundary condition: To simulate actomyosin dynamics within the contractile ring, we adapted our previously published active gel theory of the actomyosin cortex [21]. This model implements a coupled reaction-diffusion system comprising the concentrations of RhoA and actomyosin. This reaction-diffusion system is then coupled to the mechanics of an active contractile gel, representing the actomyosin network in the cortex. As described previously [12] the concentration of RhoA, *r*, is produced at some basal rate and increases its own production autocatalytically. Actomyosin, with concentration *m*, locally degrades RhoA at a rate *g*, resulting in an overall production rate of RhoA:

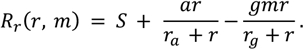

In the above equation, it is assumed that the production and degradation terms saturate with RhoA concentration. At the same time, RhoA promotes the assembly of actomyosin at a rate *k*_*a*_, which then disassembles at a rate *k*_*d*_, yielding a net production rate of actomyosin:

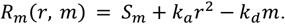

Furthermore, RhoA and actomyosin can diffuse within the cortex with diffusion constants *D*_*r*_ and *D*_*m*_, respectively, and be advected with actomyosin flows on the cortical surface. Thus, the equations describing the spatiotemporal dynamics of RhoA and actomyosin concentrations are given by:

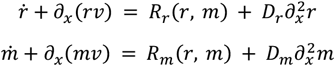

where dot denotes time derivative and *v* is the velocity of actomyosin flow. The second terms in the left-hand side of the above equations represent advection with actomyosin flows and the last terms in the right-hand side represent diffusion.

We model the actomyosin cortex as a 1-dimensional active Maxwell viscoelastic material, which behaves like a fluid at long times (due to turnover), with active stress generated by the actomyosin. The equation of motion is given by:

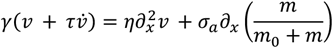

where *v* is the velocity of the cortex, *γ* is the friction between the cortex and the cytosol, η is the viscosity of the cortex, *σ*_*a*_ is the maximum active stress magnitude, with the last term representing active contraction by actomyosin which saturates at high concentrations.

The default parameters were obtained from experimental data [12]. Parameters for active stress and mechanical properties of the cortex are taken from Saha et al [107]. The diffusion rates for RhoA and actomyosin are taken from Nishikawa et al [31]. We simulate the model by numerically integrating the equations using the fipy package in python[108], using a 1-dimentional periodic box of length 140 μm.

**Table 2.**

| Default biochemical parameters | Value |
| --- | --- |
| $S_r$ | $0.01 \text{ s}^{-1}$ |
| $a$ | $0.1609 \text{ s}^{-1}$ |
| $n$ | 1 |
| $r_a$ | 0.3833 |
| $g$ | $0.1787 \text{ s}^{-1}$ |
| $r_g$ | 0.01 |
| $S_m$ | $0.0076 \text{ s}^{-1}$ |
| $k_a$ | $0.1408 \text{ s}^{-1}$ |
| $k_d$ | $0.0828 \text{ s}^{-1}$ |
| $D_r$ | $0.1 \mu\text{m}^2\text{s}^{-1}$ |
| $D_m$ | $0.01 \mu\text{m}^2\text{s}^{-1}$ |
| $\gamma_t$ | $1 \text{ Ns}^{-1} \mu\text{m}^2$ |
| $\gamma_n$ | $10 \text{ Ns}^{-1} \mu\text{m}^2$ |
| $\tau$ | 5s |
| $\lambda$ | $14.3 \mu\text{m}$ |
| $\sigma_a/\gamma$ | $49.8 \mu\text{m}^2\text{s}^{-1}$ |

The period of generated pulsatile contractions was measured using a Fourier transform of the actomyosin field at each segment of the cortex for the final 500 seconds of each simulation. The frequency power for each simulation was obtained by averaging the Fourier transform output over all spatial dimensions.

#### Simulating pulsatile RhoA activation

To apply a pulsatile stimulus with a period of 20 seconds to the excitable one-dimensional cortex, a pulsatile stimulus was simulated as the product of the positive values of 2 phase shifted sine waves with a period of 40 seconds to generate spatially alternating stimulus pulses of 20 seconds. Simulations with increasing amounts of mechanical feedback were performed for a total of 600 seconds each. The period of the observed pulsatile contraction was measured using a simple Fourier transform as described above.

#### Modeling the closure dynamics of a contractile ring

To model a closing ring, we implement our reaction-diffusion model on a series of viscoelastic edges initially arranged in a ring of radius R, connected by vertices, with positions *x*_*i*_. Edge *i* connects vertices *i* and *i* + 1 and has a tangent vector defined 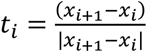, with outward normal *n*_*i*_ perpendicular to this. The edge has length *l*_*i*_ = |*x*_*i*+1_ − *x*_*i*_|.

Each edge has RhoA concentration *r*_*i*_ and actomyosin concentration *m*_*i*_ which evolve as

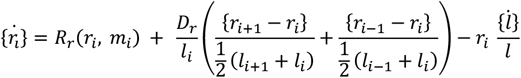

and

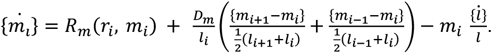

In each equation, the first term represents the production and degradation rates of RhoA and actomyosin. The second term represents diffusion across neighboring edges, which slows as the center-to-center distance between edges, e.g.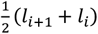, increases. The final term represents the dilution effect due to an expanding or contracting edge. Note that there is no advection term on each edge because the edges themselves move in space.

For the mechanical component, each edge has an active tension 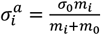 which applies contractile forces to both of the edge’s vertices, and which increases with actomyosin up to a maximum value of *σ*_0_. Each edge also acts as a spring, with rest length 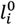 and Young’s modulus *E* producing an elastic stress 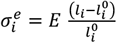. The springs are also viscoelastic, with the rest length remodeling over time 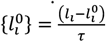 The resulting force on vertex *i* is then

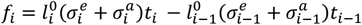

which is the sum of forces from the two edges connected to the vertex.

Finally, we update the vertex positions through force balance, in which friction balances the active and elastic forces from the contractile ring. We apply an anisotropic friction, such that friction perpendicular to the ring *γ*_*n*_ is higher than friction tangential to the ring *γ*_*t*_ as this gives our simulation more similar results to the experiments. This additional friction could be due to resistance from the cell, for example, moving the ring tangentially doesn’t deform the surface of the dividing embryo, while radially contracting the ring does, and so likely has an energy penalty associated with it. Our equation of the motion for each vertex is then

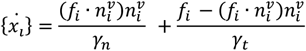

where 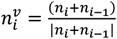 is the vertex normal, and we are projecting the force into normal and tangential components.

Within one simulation, we initially fix the radial position of the vertices while allowing for tangential motion and allow the system to evolve for 300s which allows the uniform pulses observed at the start of simulations (Fig. 6A) to develop into waves of RhoA. After this period radial motion is allowed again.

## Figures and statistical analysis

Figures were generated using Adobe Illustrator, Python, Microsoft Excel and MATLAB. Beeswarm plots (Figures 3F, 4C-D, 5B and 6D) were generated in Matlab adapting code from [109]. Wind Weighted histograms (Figure 2C, 3E-F, 4E, 5C and Supplemental Figure 2A-B [105]) were generated from Matlab scripts from the Matlab central File exchange. Statistical significance was determined using a two-tailed Student’s t test or one way ANOVA followed by a Dunnet test for multiple comparison with the same control using corresponding Matlab functions. A p value of less than 0.05 was considered significant. Respective P-values are shown in all figures. All error bars represent standard deviations. Sample size (n) are indicated in the text or the figure legend.

## Figure legends

**Supplemental Figure 1:**
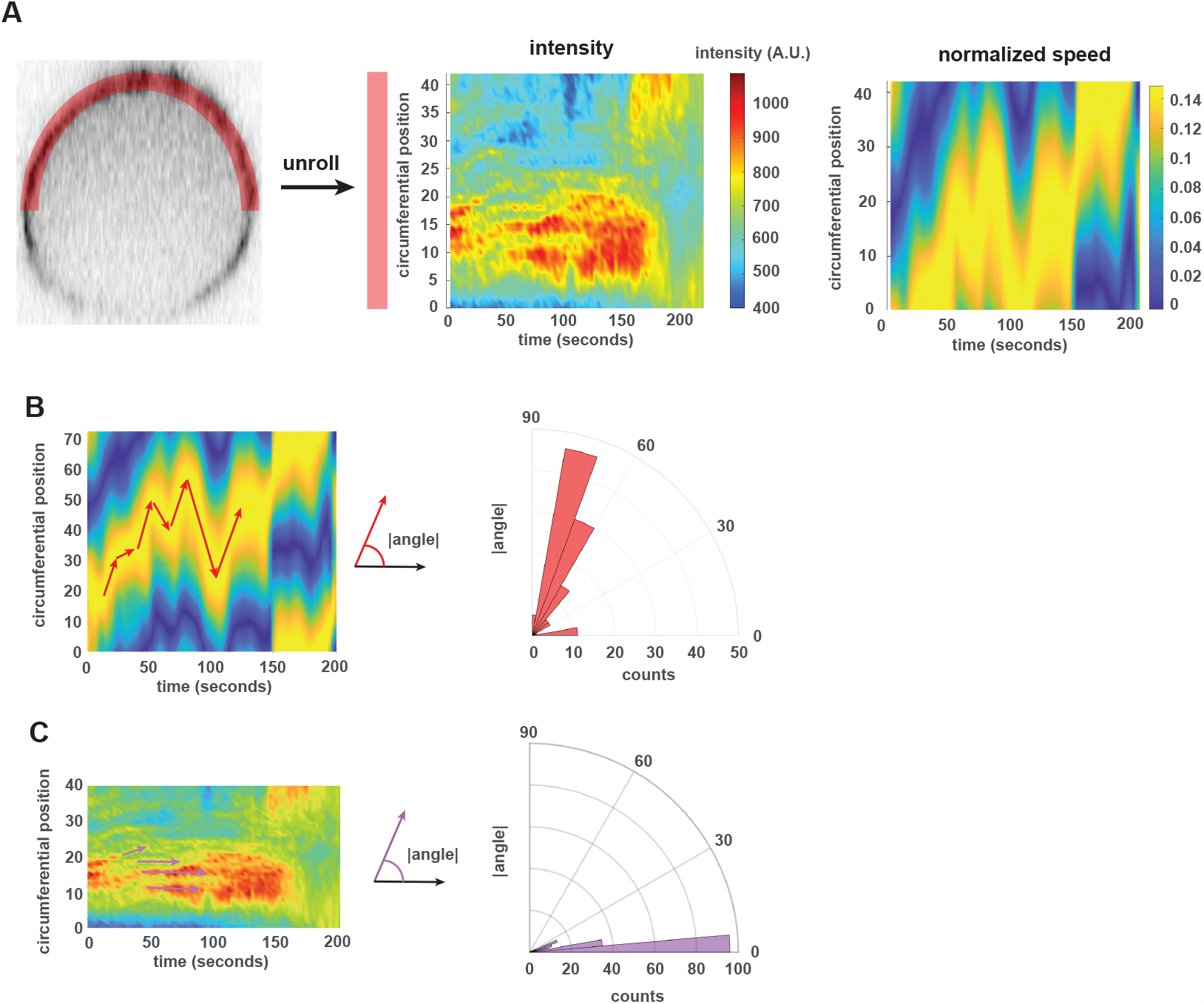
Local heterogeneities of fluorescently-tagged ANI-1 do not travel circumferentially. (A) Intensity values from only the 42 segments closest to the coverslip are plotted (left and middle) and compared to the corresponding normalized speed values (right). Intensity values are calculated as the mean of the intensity at the segment coordinate pixel and the eight pixels surrounding this position. Speed is normalized by timepoint to highlight the circumferential travel of fastest-ingressing segments. (B) Quantification of the angle of travel (left) and corresponding rose plots (right) of the fastest segment (B) and ANI-1 foci (C). Absolute angular values are shown in the plots.

**Supplemental Figure 2:**
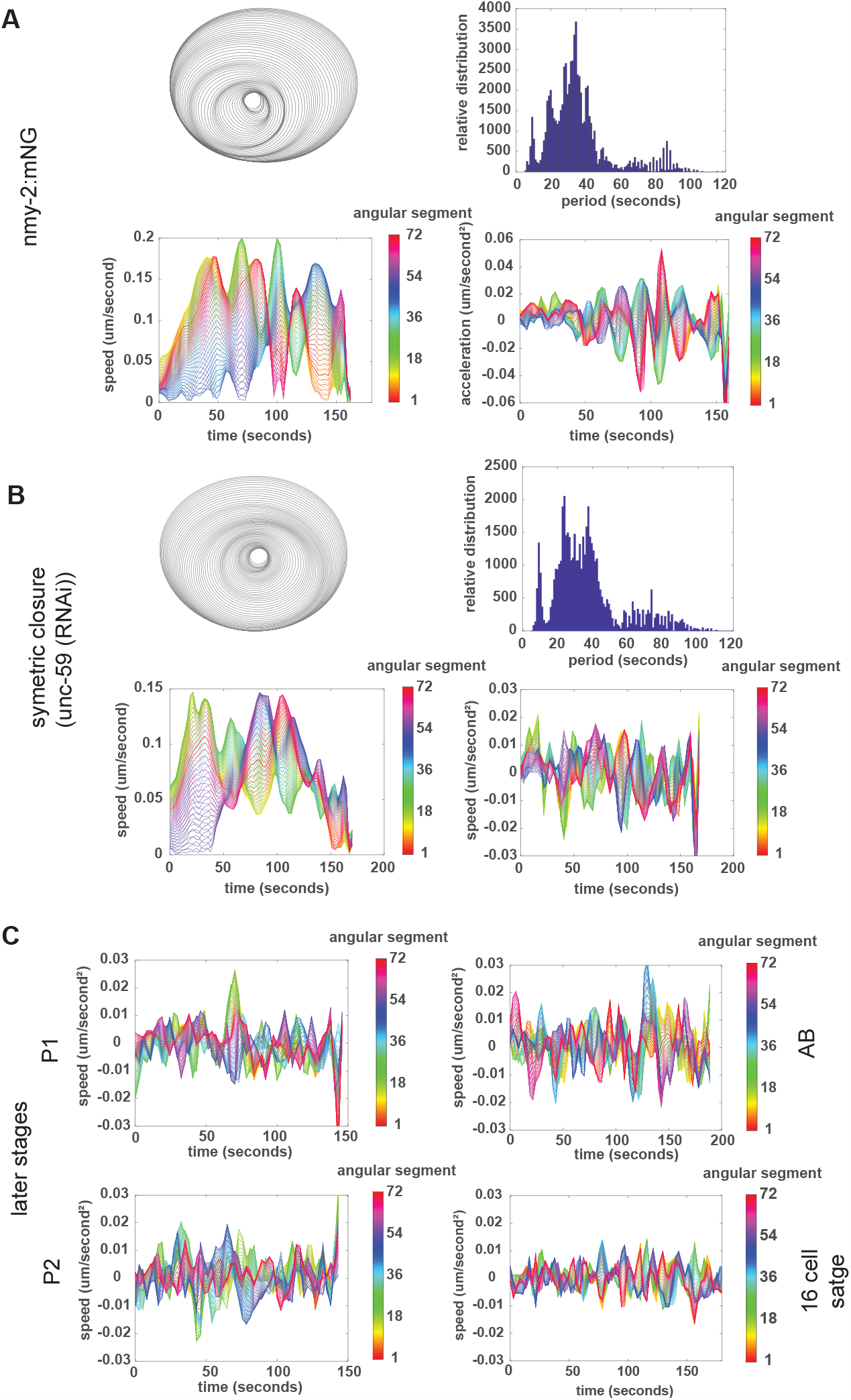
Speed oscillations are also observed in cytokinetic rings in a variety of contexts. (A) Results from analyzing ingression speed in zygotes expressing NMY-2::mNeonGreen as a ring marker (n=9). (B) Results from quantifying ingression speed in embryos depleted of UNC-59 by RNAi that ingress symmetrically (n=9). In (A) and (B), polygon fits (top left), plots of segment velocity (bottom left) and segment acceleration (bottom right) are shown for all segments of a representative embryo. Period distributions for all embryos are shown in the top right (n=9). (C) Acceleration data from a representative P1 (top left), P2 (bottom left), AB (top right), and a blastomere at the 16 cell stage (bottom right).

**Supplemental movie 1: Tracking ring closure of 72 ring segments**. Inverted contrast movie (top left) overlay of 72 angular segments (top right), coordinates of angular segments (bottom left) and cumulative angular segments (bottom right). Total movie length: 200 seconds, at 10 frames per second with a time resolution of 2.7 seconds per frame.

**Supplemental movie 2: Tracking local ingression speed during ring closure**. Inverted contrast movie (top left) overlay of 72 angular segments (top right), coordinates of angular segments (bottom left) and cumulative angular segments (bottom right). Individual dots correspond to angular segments. Colormap: ingression speed normalized for each time point. Total movie length: 200 seconds, at 10 frames per second with a time resolution of 2.7 seconds per frame.

**Supplemental movie 3. Animation of simulated ring closure data**. Colormap: ingression speed normalized for each time point. Total movie length 200 seconds of simulated time, at 27 frames per second with a time resolution of 1 second per frame.

